# Long-read single-cell RNA sequencing enables the study of cancer subclone-specific genotype and phenotype in chronic lymphocytic leukemia

**DOI:** 10.1101/2024.03.15.585298

**Authors:** Gage S. Black, Xiaomeng Huang, Yi Qiao, Philip Moos, Deepa Sampath, Deborah M. Stephens, Jennifer A. Woyach, Gabor T. Marth

**Affiliations:** Department of Human Genetics, University of Utah, Salt Lake City, UT; Department of Pharmacology and Toxicology, University of Utah, Salt Lake City, UT; Department of Hematopoietic Biology and Malignancy, The University of Texas MD Anderson Cancer Center, Houston, TX; Division of Hematology, University of North Carolina, Chapel Hill, NC; The Ohio State University Comprehensive Cancer Center, Columbus, OH

**Author notes:** **Corresponding Author Gabor Marth**, 15 North 2030 East, Room 7410B, Salt Lake City, UT 841121. G.T.M. and J.A.W. contributed equally to this work.

## Abstract

Bruton’s tyrosine kinase (BTK) inhibitors are effective for the treatment of chronic lymphocytic leukemia (CLL) due to BTK’s role in B cell survival and proliferation. Treatment resistance is most commonly caused by the emergence of the hallmark *BTK*^C481S^ mutation that inhibits drug binding. In this study, we aimed to investigate whether the presence of additional CLL driver mutations in cancer subclones harboring a *BTK*^C481S^ mutation accelerates subclone expansion. In addition, we sought to determine whether *BTK-*mutated subclones exhibit distinct transcriptomic behavior when compared to other cancer subclones. To achieve these goals, we employ our recently published method (Qiao et al. 2024) that combines bulk DNA sequencing and single-cell RNA sequencing (scRNA-seq) data to genotype individual cells for the presence or absence of subclone-defining mutations. While the most common approach for scRNA-seq includes short-read sequencing, transcript coverage is limited due to the vast majority of the reads being concentrated at the priming end of the transcript. Here, we utilized MAS-seq, a long-read scRNAseq technology, to substantially increase transcript coverage across the entire length of the transcripts and expand the set of informative mutations to link cells to cancer subclones in six CLL patients who acquired *BTK*^C481S^ mutations during BTK inhibitor treatment. We found that *BTK*-mutated subclones often acquire additional mutations in CLL driver genes, leading to faster subclone proliferation. When examining subclone-specific gene expression, we found that in one patient, *BTK*-mutated subclones are transcriptionally distinct from the rest of the malignant B cell population with an overexpression of CLL-relevant genes.

## Introduction

Chronic lymphocytic leukemia (CLL) is the most prevalent subtype of leukemia in adults, affecting ∼200,000 individuals in the United States, and is characterized by an overaccumulation of dysfunctional B cells (Byrd et al. 2004; Kipps et al. 2017). B cell receptor (BCR) signaling is crucial for cell survival and proliferation, becoming a prime target for therapeutic intervention (Burger and Chiorazzi 2013). The inhibition of Bruton’s tyrosine kinase (BTK), a kinase necessary for proper BCR signaling, has proven to be an effective treatment for most patients (Petro et al. 2000; Herman et al. 2011; Byrd et al. 2013; Pal Singh et al. 2018). However, secondary resistance to treatment develops in ∼20% of patients over time, leading to poor clinical outcomes, including shorter survival (Nakhoda et al. 2023). The *BTK*^C481S^ mutation is the most common cause of BTK inhibitor resistance (up to 80% of relapses) and confers drug resistance via impairing binding of the drug to CLL cancer cells, transforming the normal covalent BTK inhibitor binding to a noncovalent bond (Woyach et al. 2017; Sedlarikova et al. 2020). The evolution of mutations in additional CLL-driver genes has also been shown to contribute to treatment resistance (Landau et al. 2015; Komarova et al. 2014; Burger et al. 2016). Our present study aims to investigate whether the presence of additional CLL driver mutations in cancer subclones harboring a *BTK*^C481S^ mutation accelerates subclone expansion and whether *BTK-*mutated subclones exhibit distinct transcriptomic behavior when compared to other cancer subclones.

Previously, we have demonstrated the ability to use bulk DNA to deconvolute cancer subclone structures (Qiao et al. 2014; Brady et al. 2017; Than et al. 2018; Huang et al. 2021; Black et al. 2022), as well as using single-cell RNA sequencing (scRNA-seq) data to genotype individual cells and study subclone-specific transcriptomic behavior using scBayes (Qiao et al. 2024). We use whole-exome sequencing (WES) or whole-genome sequencing (WGS) of tumor/normal pairs to identify somatic mutations and reconstruct the subclone structure of the tumor as well as its evolution across tumor progression. Individual cells from the scRNA-seq data are assigned to genomic subclones based on the presence or absence of subclone-defining mutations. By combining subclone identity with single-cell gene expression information, this approach enables a subclone-specific gene expression analysis.

Until recently, scRNA-seq has been carried out almost exclusively with short-read sequencing technologies. A disadvantage of traditional short-read scRNA-seq is the 5’ or 3’ bias, where the vast majority of the reads are concentrated at the 5’ or 3’ ends of the transcript (Ziegenhain et al. 2017). This bias limits our ability to determine the cell’s genotype at the sites of somatic tumor mutations farther, i.e., at greater than sequencing read-length distances, from the priming site. Long-read scRNA-seq provides a promising alternative, as it can provide coverage across the full transcript length. MAS-seq, a long-read scRNA-seq solution by Pacific Biosciences (PacBio), is a ready-to-use kit that concatenates cDNAs generated from the 10X Chromium platform to create long composite molecules that can be sequenced with over 99.9% accuracy via HiFi sequencing (Al’Khafaji et al. 2023; Wenger et al. 2019). We hypothesized that the highly accurate full-transcript coverage afforded by MAS-seq would yield improved coverage of somatic mutations falling outside the standard short read-length distance of priming sites, thus enabling enhanced single-cell genotyping and subclone assignment.

In a recent study, we investigated subclonal evolution using WES in 38 patients with CLL treated with a BTK inhibitor (Supplemental Table 1) (Black et al. 2022). We found that the evolution of subclones containing mutations in CLL driver genes within the first two years of treatment had a significant association with eventual relapse. Among these patients, we identified six who developed at least one subclone harboring a *BTK*^C481S^ resistance mutation. Here, we performed long-read scRNA-seq with MAS-seq on samples from these six patients taken before BTK inhibitor treatment and at the time of relapse to use in conjunction with the WES data to study the co-occurrence of *BTK*^C481S^ and additional CLL driver mutations, as well as *BTK*-mutant subclonal phenotypes.

## Results

### Long-read sequencing with MAS-seq expands transcript and variant coverage

We generated long-read scRNA-seq data for the pre-treatment and relapse samples from the six CLL patients who developed *BTK*^C481S^ mutations during treatment. Cell isolation was performed using the 10X Genomics Chromium 3’ Single Cell Kit, followed by cDNA concatenation and HiFi sequencing with PacBio’s MAS-seq technology. Four samples were sequenced on the Sequel II system, while eight samples were sequenced on the Revio system. When comparing the number of reads produced by the two sequencing instruments, we find that the Revio produced ∼4x the number of reads per sample (Figure 1A). Samples sequenced on the Sequel II produced an average of 3,363 reads per cell, with 4,251 cells per sample (Table 1). In contrast, samples sequenced on the Revio produced an average of 9,283 reads per cell, with 6,239 cells per sample. These outcomes highlight the Revio’s superior data generation capabilities, offering a more cost-effective long-read sequencing solution and enabling a more robust analysis. All samples across both platforms maintained a high median base quality, consistently at or above Q30.

**Figure 1.**
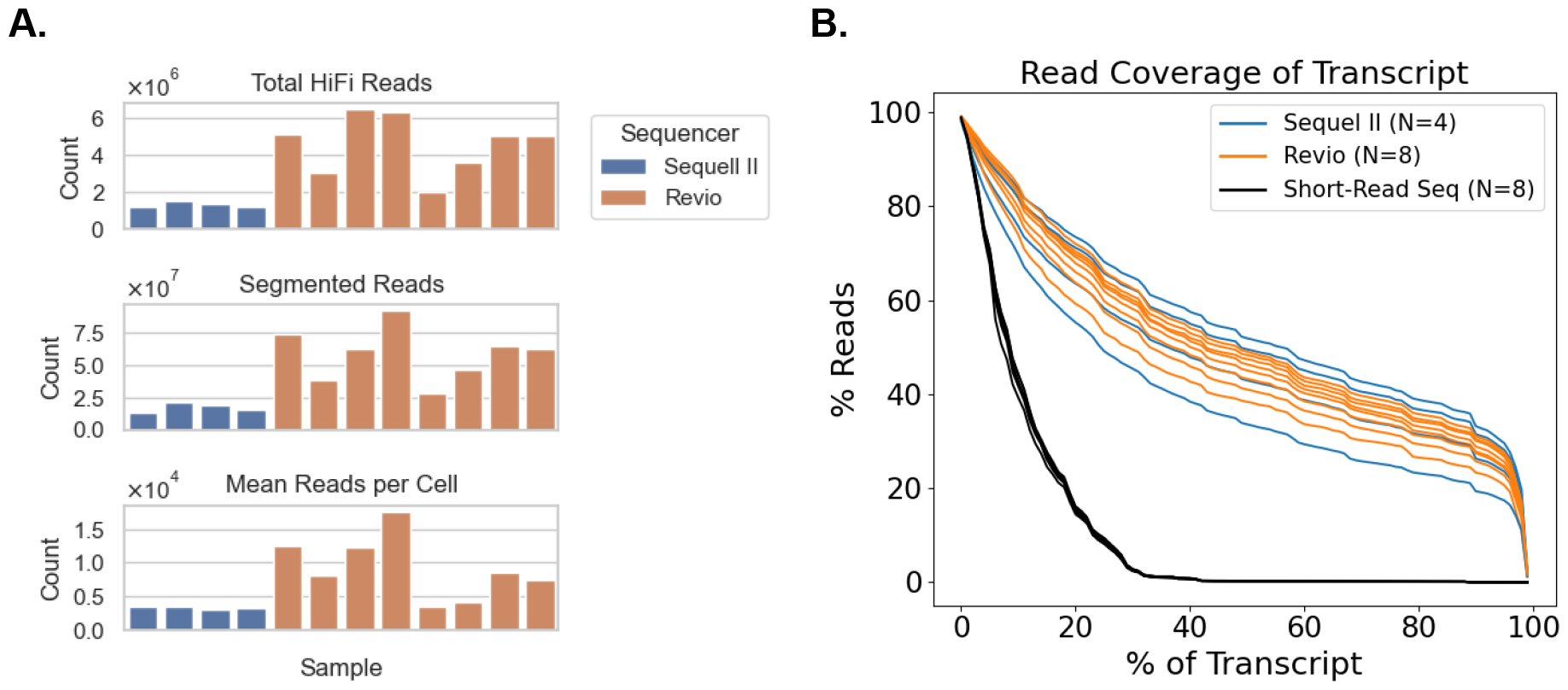
Long-read scRNA sequencing metrics. A) Comparison of the total number of HiFi reads, the total number of segmented reads, and the mean reads per cell for each sample, colored by the sequencer used. B) The canonical transcript coverage for each read aligning to a given gene is calculated for all protein-coding genes with at least one read aligned to it. The percentage of reads covering X% of the given transcript is plotted for each sample, colored by the sequencer used for the sample. Eight short-read samples are included in black for comparison.

**Table 1.**
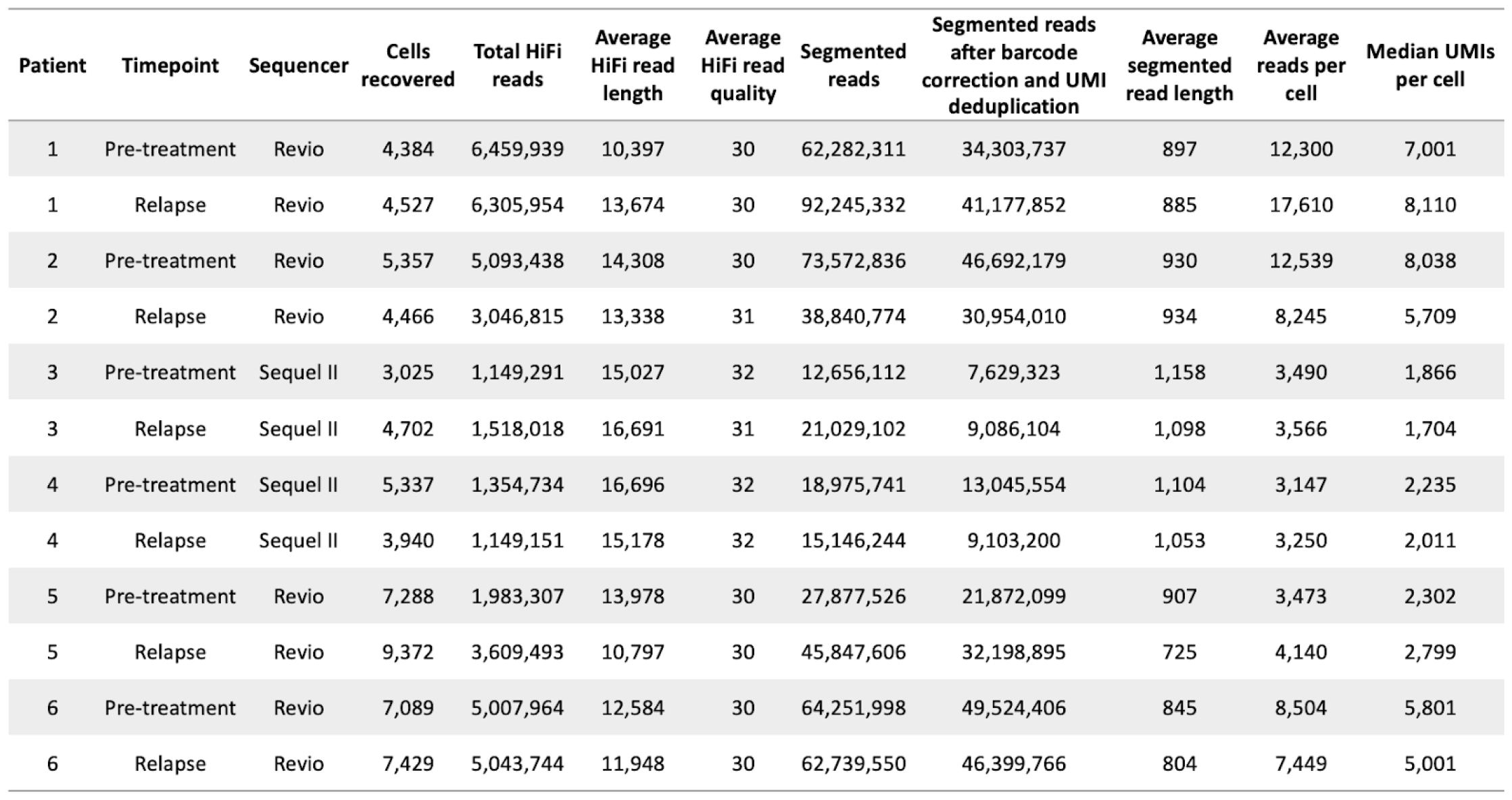
Long-read scRNA sequencing metrics for each sample across the six patients.

| Patient | Timepoint | Sequencer | Cells recovered | Total HiFi reads | Average HiFi read length | Average HiFi read quality | Segmented reads | Segmented reads after barcode correction and UMI deduplication | Average segmented read length | Average reads per cell | Median UMIs per cell |
| --- | --- | --- | --- | --- | --- | --- | --- | --- | --- | --- | --- |
| 1 | Pre-treatment | Revio | 4,384 | 6,459,939 | 10,397 | 30 | 62,282,311 | 34,303,737 | 897 | 12,300 | 7,001 |
| 1 | Relapse | Revio | 4,527 | 6,305,954 | 13,674 | 30 | 92,245,332 | 41,177,852 | 885 | 17,610 | 8,110 |
| 2 | Pre-treatment | Revio | 5,357 | 5,093,438 | 14,308 | 30 | 73,572,836 | 46,692,179 | 930 | 12,539 | 8,038 |
| 2 | Relapse | Revio | 4,466 | 3,046,815 | 13,338 | 31 | 38,840,774 | 30,954,010 | 934 | 8,245 | 5,709 |
| 3 | Pre-treatment | Sequel II | 3,025 | 1,149,291 | 15,027 | 32 | 12,656,112 | 7,629,323 | 1,158 | 3,490 | 1,866 |
| 3 | Relapse | Sequel II | 4,702 | 1,518,018 | 16,691 | 31 | 21,029,102 | 9,086,104 | 1,098 | 3,566 | 1,704 |
| 4 | Pre-treatment | Sequel II | 5,337 | 1,354,734 | 16,696 | 32 | 18,975,741 | 13,045,554 | 1,104 | 3,147 | 2,235 |
| 4 | Relapse | Sequel II | 3,940 | 1,149,151 | 15,178 | 32 | 15,146,244 | 9,103,200 | 1,053 | 3,250 | 2,011 |
| 5 | Pre-treatment | Revio | 7,288 | 1,983,307 | 13,978 | 30 | 27,877,526 | 21,872,099 | 907 | 3,473 | 2,302 |
| 5 | Relapse | Revio | 9,372 | 3,609,493 | 10,797 | 30 | 45,847,606 | 32,198,895 | 725 | 4,140 | 2,799 |
| 6 | Pre-treatment | Revio | 7,089 | 5,007,964 | 12,584 | 30 | 64,251,998 | 49,524,406 | 845 | 8,504 | 5,801 |
| 6 | Relapse | Revio | 7,429 | 5,043,744 | 11,948 | 30 | 62,739,550 | 46,399,766 | 804 | 7,449 | 5,001 |

To examine the transcript coverage afforded by reads generated from MAS-seq, we selected the Ensembl canonical transcripts for all protein-coding genes and calculated the percentage of transcript coverage provided by each read mapping to the given gene (Figure 1B). In addition, we calculated this same coverage within pre-treatment and relapse samples from four patients (within our original 38-patient cohort but not among the six patients in the present study) that were sequenced using short-read scRNA-seq. We find comparable transcript coverage across all samples sequenced with MAS-seq, with 50% of reads covering at least 41.3% of the transcript length and 32.5% of reads covering >80% of the canonical transcript on average. In comparison, we found that the short-read sequencing samples had 50% of reads covering at least 9.1% of the transcript length and only 0.1% of reads covering >80% of the canonical transcript on average. This comparison illustrates the significant increase in transcript coverage provided by each read when using MAS-seq.

To determine our ability to identify multiple mutations within a given cell, we selected heterozygous germline variants from each sample and calculated the percentage of variants with coverage in each cell (Supplemental Figure 1). Figure 2A illustrates these coverage metrics by comparing the sample with the best coverage from the Sequel II, Revio, and short-read sequencing technologies. The sample sequenced on the Sequel II had 12.61% of heterozygous germline variants covered by RNA-seq data in 1% or more of the cells and 1.73% of variants covered in 10% or more of cells (13,403 total variants). The sample sequenced on the Revio had a clear increase in coverage, with 18.59% of variants covered in 1% or more of the cells and 3.85% of variants covered in 10% or more of cells (13,709 total variants). In comparison, the sample sequenced with short-reads had 7.61% of variants covered in 1% or more of the cells and 1.59% of variants covered in 10% or more of cells (12,954 total variants). Furthermore, we characterized each heterozygous germline variant by its distance from the priming end of the transcript to determine the impact of this distance on variant coverage (Figure 2B, Supplemental Figure 2). Samples sequenced with MAS-seq on either the Revio or Sequel II show uniform variant coverage across all positions in the transcript. In contrast, the short-read sequencing shows a bias in variant coverage for those that are <400 base pairs (bp) from the priming site. These results indicate that the Revio system provides meaningfully increased variant coverage per cell compared to short-read sequencing and that this coverage can be seen throughout the entire transcript. The increased, uniform coverage afforded by MAS-seq improves our ability to genotype cells at multiple subclone-defining somatic mutation sites in a single cell, enhancing the quality of cell assignments to cancer subclones.

**Figure 2.**
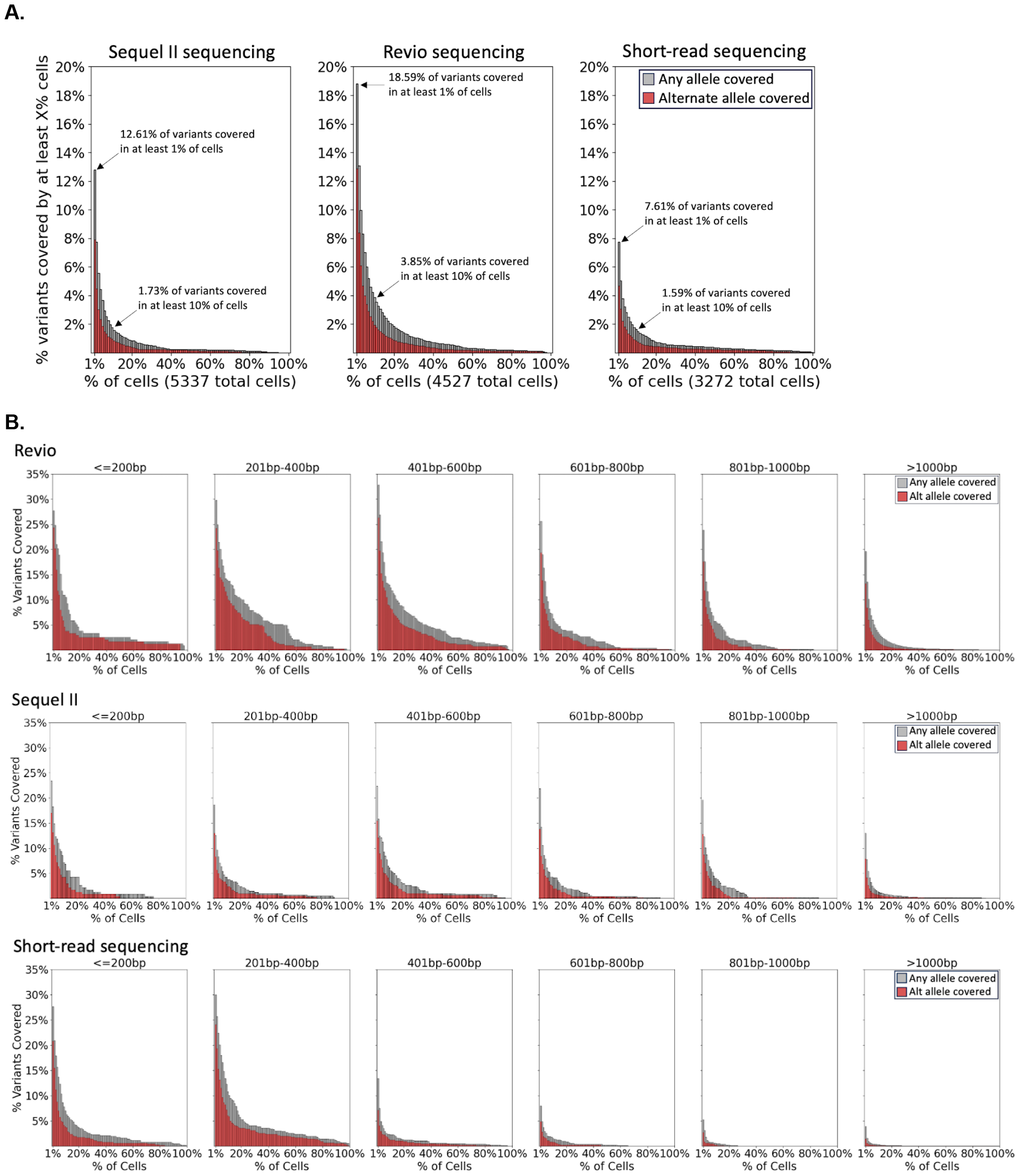
Variant coverage provided by each scRNA-seq technology. A) The overall variant coverage provided by the Sequel II and Revio compared to Illumina short-reads. The percentage of cells covering each heterozygous germline variant in the patient’s WES data is used to determine the percent of variants covered by at least X% of cells. B) The variant coverage binned by the variant’s distance from the priming site, as indicated above each plot (bp = base pairs).

### Full-transcript coverage allows for the identification and visualization of single-cell genotypes

We have recently published a novel computational method, scBayes, to study cancer subclone-specific expression phenotypes by combining scRNA-seq and bulk DNA sequencing-based subclone structures (Figure 3A) (Qiao et al. 2024). This approach enables genotype inference for mutations lacking sequencing coverage when genotypes are present for other subclone-specific mutations within the same subclone. The cancer subclone structures of the six patients in this study have been extensively characterized in our previous work (Black et al. 2022), allowing us to apply scBayes and assign each cell to a clone of origin. Briefly, scBayes genotypes each cell by examining the presence of subclone-defining somatic mutations (as discovered from bulk DNA data, Figure 3B, y-axis) in the scRNA-seq reads of the cell (Figure 3B, x-axis) and uses a Bayesian probabilistic framework to identify the most likely clone of origin (Figure 3B, x-axis color bars) for the cell. We use a scatterplot-like visualization (Figure 3B) to illustrate the cell genotypes, showing whether the mutant allele was observed (red) for each variant in each cell, if only the reference allele was observed (green), or if there was no read coverage for the variant (white). Once cells are assigned, those of the same subclone can be grouped together and compared to cells of other subclones for subclone-specific gene expression analysis (Figure 3C). Long-read RNA sequencing is particularly suitable for such an approach as it maximizes the chance that any somatic mutation is covered by sequencing.

**Figure 3.**
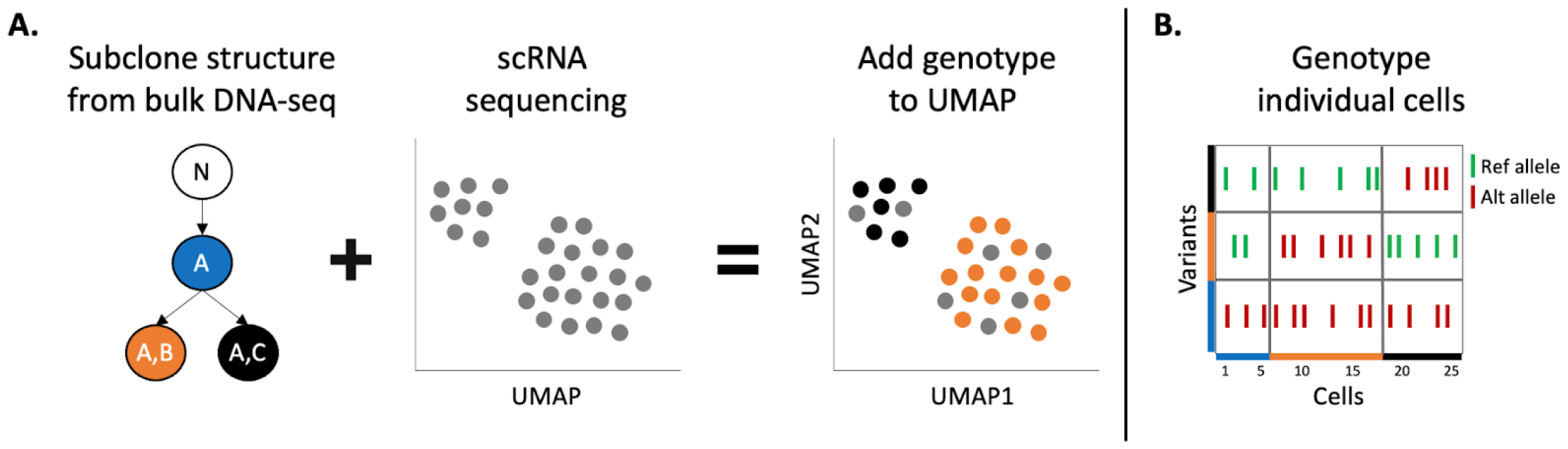
Overview of workflow to identify and use cell genotypes. A) Pre-determined subclone structures with accompanying somatic variants are used to genotype individual cells in scRNA-seq data. Cells are assigned to a pre-determined subclone based on the presence or absence of subclone-defining mutations. Subclone assignments are then used to group cells of the same subclone to identify subclone-specific gene expression. B) Genotype matrix plots visualize the genotypes of all cells at each variant of interest, showing green for reference allele, red for alternate allele, and white for no coverage.

### Long-read scRNA-seq enables confirmation or refinement of subclone structures

To determine whether the structures previously identified through the WES data were corroborated by the long-read scRNA-seq data, we analyzed the subclone structure of each patient at a single-cell level. In five of the six patients, the subclone structure identified in the WES data was confirmed by the long-read scRNA-seq data (Figure 4, Supplemental Figure 3, and Supplemental Figure 4). We illustrate such concordance in Patient 1, where the WES data showed a linear pattern of subclonal evolution (Figure 4A). To visualize this same pattern of evolution within the scRNA-seq samples, we created a genotype matrix plot showing each cell’s genotype at each somatic variant within the sample. Figure 4B shows a representation of the genotype matrix plot for the two samples from this patient. Cells belonging to the same subclone show similar genotype patterns, seen as clusters of red within the plot. The pretreatment sample depicts five distinct subclones, each harboring the variants of the previous subclone. In the relapse sample, we again see the linear pattern of evolution where the *BTK*-mutated subclone (SC6) emerged and became the dominant subclone. By genotyping cells in the long-read scRNA-seq data, we conclude that the subclone structure identified through WES accurately represented the true subclonal heterogeneity in this patient’s cancer.

**Figure 4.**
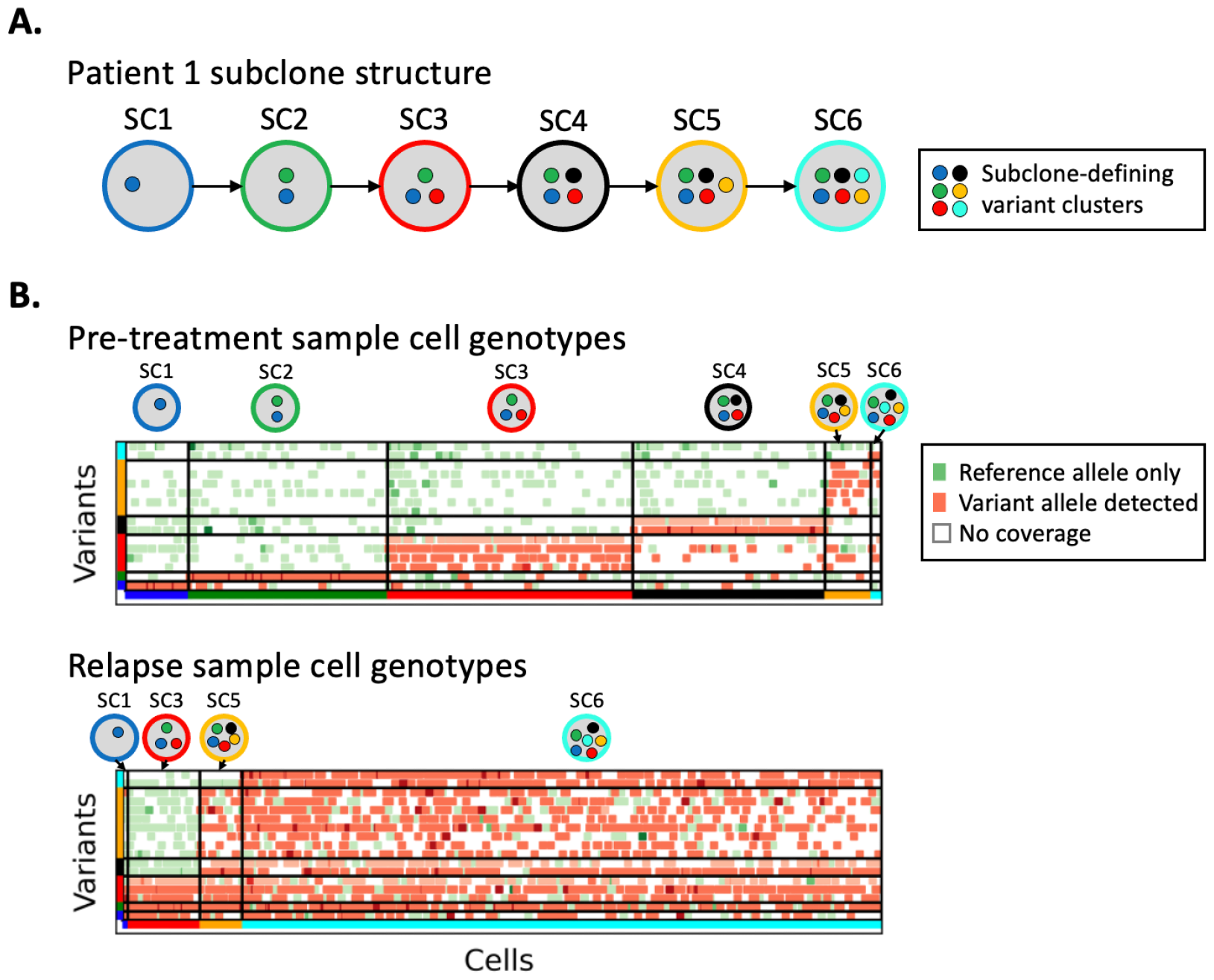
Visualization of single-cell genotypes to identify subclone structures. A) The subclone structure of Patient 1 identified in the bulk DNA sequencing data. Subclones are depicted by the colored circles, with representative variant clusters inside each circle. B) The cell genotypes at subclone-defining variants in Patient 1, with green markers representing only reference alleles present in the scRNA-seq reads at the given variant location within the cell and red markers indicating at least one scRNA-seq read in the cell contains the somatic variant allele. Darker marker coloring indicates an increased number of reads supporting that genotype. Variants and cells are grouped by their subclone assignment.

For the remaining patient (Patient 3, see Figure 5), the resolution provided by long-read scRNA-seq highlighted the need for further refinement of the subclone structure that was previously constructed. In the WES of Patient 3, two *BTK*^C481S^ mutations were identified at two different bases of the same codon (*BTK* c.1543T>A and *BTK* c.1544G>C), exhibiting similar variant allele frequencies (VAFs) in the samples where they were detected (Figure 5A). Because VAFs are used to cluster variants and identify subclones in bulk DNA sequencing, these *BTK* mutations were initially clustered into the same subclone (SC2) alongside other variants with matching VAFs. We utilized the long-read scRNA-seq data to investigate individual cells with coverage at these *BTK* mutation sites and found that these two *BTK* mutations were, in fact, part of two distinct subclones. The genotype matrix plot for the relapse sample of this patient shows that cells with the *BTK* c.1544G>C mutation and its associated mutations never co-exist with the *BTK* c.1543T>A mutation (Figure 5B). This increased resolution of the subclonal architecture afforded by the long-read scRNA-seq data allows us to conclude that rather than being on different haplotypes within the cells of a single subclone, these mutations belong to two distinct *BTK*-mutated subclones that had similar cellular prevalences in the patient sample (Figure 5C).

**Figure 5.**
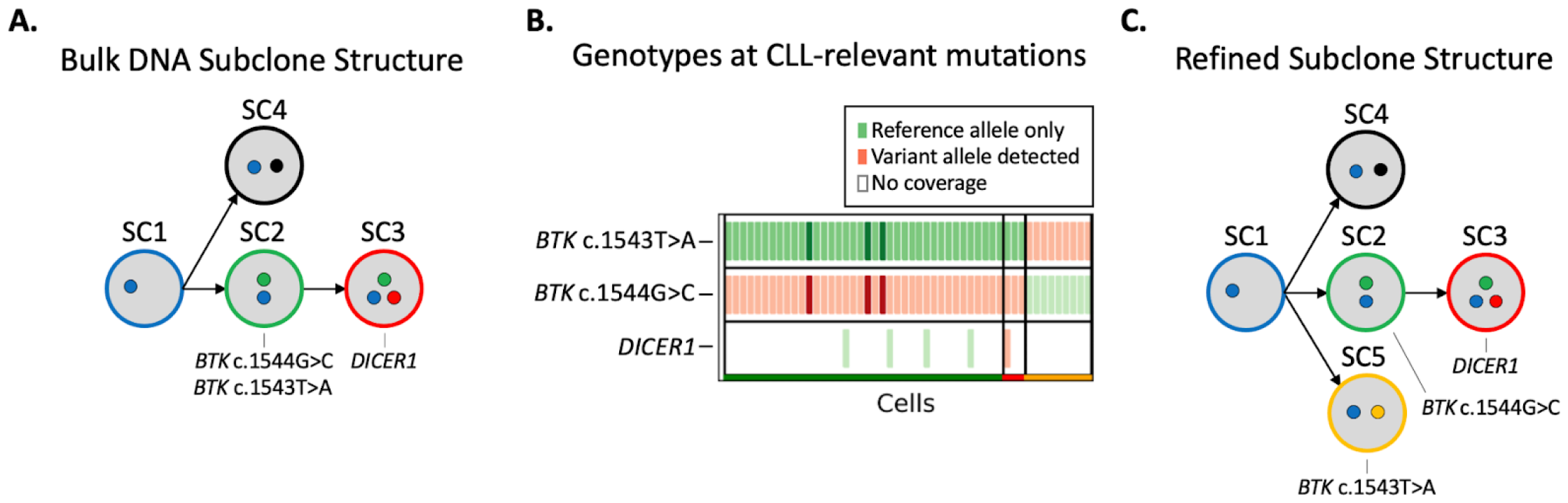
Refining the subclone structure of Patient 3. A) The subclone structure identified in the bulk DNA sequencing data. CLL-relevant gene mutations are annotated under the subclone they are found in. B) The genotype matrix plot from the relapse sample of Patient 3 enables refinement of the original subclone structure. Green markers indicate that only reference alleles were present in the scRNA-seq reads at the given variant location within the cell, and red markers indicate that at least one scRNA-seq read in the cell contains the somatic variant allele. Darker coloring indicates an increased number of reads supporting that genotype. Only the CLL-relevant mutations are included for increased resolution to differentiate subclones. C) The refined subclone structure that depicts the subclone containing the *BTK* c.1543T>A mutation is independent of the subclone containing the *BTK* c.1544G>C and *DICER1* mutations.

### The ability to identify variants co-occurring in the same tumor cells allowed us to identify *BTK*-mutated cells with additional CLL driver gene mutations

Next, we wanted to determine whether cells with *BTK*^C481S^ mutations contained additional mutations in CLL driver genes. For this analysis, we genotyped *BTK*-mutated cells to identify the presence of mutated alleles in CLL-driver genes. We found that the *BTK*-mutated subclones of all six patients contained additional mutations in driver genes at the time of relapse (Table 2). In all but one patient, these mutations were present in the cells before the *BTK* mutation developed and were detectable in the pretreatment sample (Supplemental Figure 3).

**Table 2.** The co-occurrence of mutations in CLL driver genes in *BTK*-mutated subclones.

| Patient ID | CLL mutations co-occurring with *BTK*<sup>C481S</sup> | CLL mutations were present before *BTK*<sup>C481S</sup> developed |
| --- | --- | --- |
| 1 | *ASXL1*, *SF3B1*, *KMT2D*, *CREBBP* | Yes |
| 2 | *BRAF*, *SAMHD1*, *BCOR*, *NFKBIE*, *EGR2* | Yes |
| 3 | *DICER1* | No |
| 4 | *TP53* | Yes |
| 5 | *NOTCH1* | Yes |
| 6 | *SF3B1*, *POT1* | Yes |

Patient 3, who had two independent *BTK*-mutated subclones emerge with similar allele frequencies, was the exception. When each of these *BTK*-mutated subclones developed, neither had any additional CLL-associated mutations that were detectable (Supplemental Figure 4). One of these subclones continued to evolve, developing a mutation in *DICER1*—a gene recently implicated in CLL (Knisbacher et al., Nature Genetics, 2022). At the time of relapse, the *DICER1* and *BTK* co-mutated subclone demonstrated a higher prevalence in the sample compared to the *BTK*-only subclone.

Patient 5 also developed two independent *BTK*-mutated subclones during treatment (Supplemental Figure 4). The first to develop did not harbor any additional CLL mutations. However, the second *BTK*-mutated subclone arose from a cell population containing a *NOTCH1* mutation, a known CLL driver gene (Arruga et al. 2014; Fabbri et al. 2011). This second *BTK*-mutated subclone rapidly expanded and became the dominant subclone in the relapse sample. This data shows that subclones with co-occurring mutations in *BTK* and additional CLL driver genes expanded more rapidly than subclones with just *BTK* mutations alone.

### Subclone assignments enabled by long-read scRNA-seq reveal transcriptomically distinct

#### *BTK*-mutated subclones

After subclone assignment, cells underwent further analysis with Seurat, a standard tool for single-cell gene expression clustering (Hao et al. 2021). Here, our goal was to determine whether subclone-level gene expression patterns could be identified within the long-read scRNA-seq data. Normalized gene counts were used to cluster cells based on gene expression patterns. Utilizing the cell barcodes from the scBayes subclone assignments, we overlaid subclone identities onto these gene expression-based cell clusters, facilitating inter-subclone gene expression comparisons.

In the relapse sample of Patient 1, we identified two distinct gene expression clusters. Mapping subclone assignments onto these clusters showed that the larger cluster was primarily composed of cells from the *BTK*-mutated subclone, which had emerged as the dominant clone in that sample (Figure 6A). The smaller cluster consisted of cells from an earlier ancestral subclone, highlighting the transcriptional divergence between these two populations. Of the 6 patients, Patient 1 was the only one where the BTK-mutated subclone created a distinct cluster of cells within the transcriptomic data (Supplemental Figure 5)

**Figure 6.**
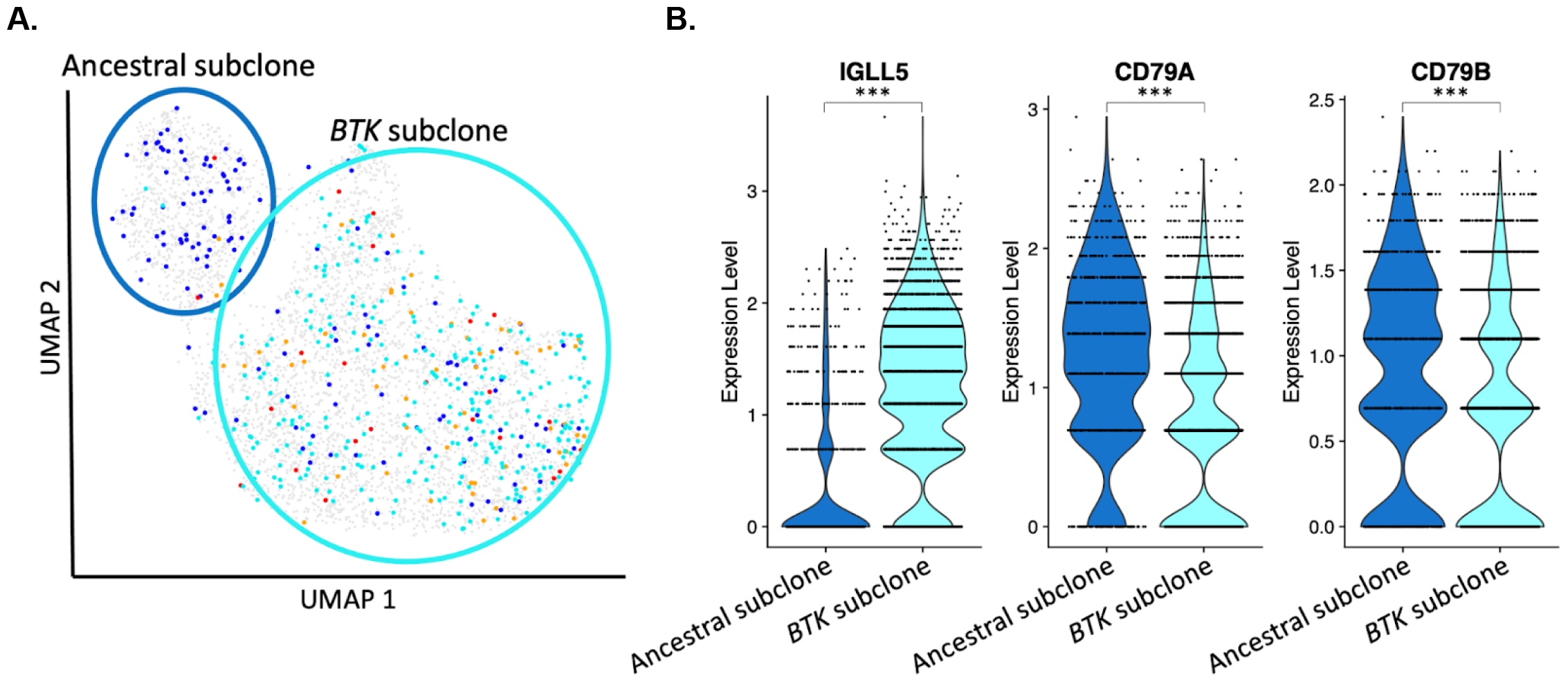
Using cell assignments to identify subclone-specific gene expression patterns. A) Mapping subclone assignment to clustered cells enables the identification of phenotypically distinct subclones. B) Differential gene expression analysis between subclones illuminates over- and under-expressed genes within the *BTK*-mutated subclone. (***) adjusted p-value < 0.001.

Differential gene expression analysis between the two clusters in Patient 1 found that *IGLL5*, a gene implicated in CLL (Kasar et al. 2015; Pérez-Carretero et al. 2020; Deng et al. 2023), was overexpressed in the *BTK*-mutated subclone when compared to the ancestral subclone (Figure 6B). *CD79A* and *CD79B*, two genes upstream of *BTK* in the B cell receptor pathway, show lower levels of expression in the *BTK*-mutated subclone when compared to the ancestral subclone. These findings highlight the dual utility of long-read scRNA-seq in elucidating both genomic and transcriptomic subclonal dynamics within a single assay.

## Discussion

This study utilizes MAS-seq, PacBio’s long-read scRNA-seq technology, to comprehensively analyze the subclonal dynamics and gene expression patterns within CLL, offering insights into the cellular heterogeneity and mechanisms underlying BTK inhibitor treatment resistance. This technological advancement represents an improvement over short-read scRNA-seq, primarily by improving on its limitations in transcript coverage and mutation site resolution. By comparing samples sequenced on the Sequel II and Revio instruments, we see a significant improvement in the quantity of data provided by the latter. When comparing to samples sequenced using traditional short-read scRNA-seq, we see increased transcript and variant coverage provided by MAS-seq. Despite these improvements, MAS-seq is still limited by the number of variants with coverage within a single cell. This limitation in variant coverage could be attributed to technical factors, such as a relative scarcity of reads from certain cells, or biological factors, such as lower expression levels of the genes of interest compared to those with higher expression, which consequently receive more sequencing reads. To address this limitation, strategies like CRISPR/Cas9-mediated depletion of over-represented cDNAs, e.g., those belonging to long-noncoding RNA, can be implemented to enhance the proportion of informative reads (Pandey et al. 2022; Wang and Adler 2023). Alternatively, targeted enrichment strategies, such as hybridization capture for gene panels, could be utilized to enhance the coverage of key genes, improving the detection of variants (Pokhilko et al. 2021). These approaches would increase the read depth at variants of interest, improving cell genotyping and gene expression analysis.

Our approach provides a framework for utilizing long-read scRNA-seq to identify patterns of subclonal evolution within a patient sample. When used in conjunction with bulk DNA-seq data, subclones can be identified and refined with greater resolution than using bulk DNA alone. In one patient, we found that the data provided by the bulk DNA-seq was insufficient to identify the correct subclone structure due to two different subclones having very similar cellular prevalences. By interrogating the genotypes within the cells of these subclones, we could determine that they were two independent subclones, and rectify the bulk DNA-based subclone structure.

When investigating the transcriptomic behavior within the patient samples, we identified one patient where the *BTK*-mutated subclone represents the cells of a distinct gene expression cluster. This separation enables an inter-subclone differential gene expression analysis to identify genes that were over- or under-expressed in the *BTK*-mutated subclone. No other patients showed the same pattern of *BTK*-mutated subclones being responsible for an isolated cluster within the transcriptomic data. Increasing the number of reads per cell would allow for a more robust transcriptomic analysis within these cell populations. To improve the quantification of gene expression while maintaining the additional information provided by long-read scRNA-seq, sequencing libraries could also be sequenced using a short-read technology to increase the number of reads per gene (Torre et al. 2023; Mincarelli et al. 2023). While these reads will not provide the same full-transcript coverage, they will enable a higher-resolution gene expression analysis within the sample.

Two of the six patients included in this study developed two independent *BTK*-mutated subclones. We found that subclones harboring both a *BTK*^C481S^ mutation and additional mutations in CLL driver genes have increased expansion compared to those without the additional driver mutations at the time of relapse. These findings suggest that the co-occurrence of these mutations may confer an increased cell survival advantage and contribute to resistance to therapy. Understanding the *BTK*-mutated subclones’ dynamics can provide valuable insight into the pattern of symptoms that occur along with clinical relapse and may help to determine how quickly relapse will occur. Understanding subclonal evolution can help detect resistant clones early and guide specific subclone-directed therapy to reduce further selection of aggressive subclones. Combination therapy may be required to treat CLL with multiple subclones. Alternatively, the presence of multiple subclones may result in a clinical indication to shift to alternate therapies with distinct mechanisms of action from the current therapy.

## Methods

### CLL Patient Cohort

We received samples from 6 CLL patients treated with BTK inhibitors at The Ohio State University to undergo long-read scRNA-seq. Blood samples and other standard clinical data were collected through an IRB-approved tissue procurement protocol during routine clinical visits.

### B cell isolation and library preparation

Peripheral blood mononuclear cells (PBMCs) were initially isolated from whole blood using ficoll density gradient centrifugation. PBMCs were then viably frozen. Prior to sequencing, samples were processed by sequentially using the EasySep™ Dead Cell Removal (Annexin V) Kit (cat # 17899), followed by buffer exchange and B cell selection using the EasySep™ Human B Cell Enrichment Kit II Without CD43 Depletion (cat # 17963) from Stemcell Technologies (Vancouver, BC). Following B cell selection, the 10X Genomics Chromium Next GEM Single Cell 3’ Reagent Kits v3.1 (Dual Index) was used for cell barcoding and cDNA generation. The standard protocol was followed until the end of step 2, stopping before cDNA cleavage. The resulting cDNA was used as input for the MAS-Seq for 10x Single Cell 3’ kit by PacBio.

### Long-read sequencing and data processing

Four samples (pre-treatment and relapse samples from patients 3 and 4) were sequenced on the PacBio Sequel II system using 8M SMRTCells for 30 hours. The remaining samples were sequenced on the PacBio Revio system using Revio SMRT Cells for 24 hours. Raw HiFi reads were segmented into representative cDNAs with SMRT Link v12.0.0.177059 using the 10X Chromium single cell 3’ cDNA primers barcode set and MAS-Seq Adapter v1 (MAS16) barcode set. We used PacBio’s Iso-Seq single-cell workflow to trim, tag, and align reads using the default parameters of the tools distributed via Bioconda (https://github.com/PacificBiosciences/pbbioconda). Briefly, Lima v2.7.1 removed primers from the segmented reads. Isoseq v4.0.0 added tags for unique molecular identifiers (UMIs) and barcodes, followed by trimming of the PolyA and primer sequences. Isoseq was then used to correct cell barcodes and deduplicate reads. Finally, pbmm2 v1.10.0 aligned reads to the reference genome using the GRCh38 reference sequence. Pigeon v1.2.0 generated Seurat-compatible files for downstream transcriptomic analysis.

### Reconstructing subclone structures from whole-exome sequencing data

In a previous study, we collected 200X whole-exome sequencing data for each patient at multiple timepoints throughout their treatment (Black et al. 2022). Reads were aligned to the GRCh38 reference genome with BWA-mem v0.7.17 (Li and Durbin 2009). Samblaster (Faust and Hall 2014) removed duplicate reads from the output generated by the alignment. Samtools v1.16 (Li et al. 2009) was used to merge and sort BAM files by leftmost coordinates. All data processing commands were run using the default parameters for each tool.

We used Freebayes v1.3.4 (Garrison and Marth 2012) to call variants within each patient. A minimum alternate fraction filter of 0.5 was used along with the following options: allele-balance-priors-off, report-genotype-likelihood-max, genotype-qualities, pooled-discrete, and pooled-continuous. The VCF that was generated from this script represents all variants within the patient and their allele count at each timepoint during BTK inhibitor treatment. Using Bedtools intersect v2.28.0 (Quinlan and Hall 2010), we removed any variants that did not fall within exonic regions or accessible regions specified by the 1000 Genomes project (1000 Genomes Project Consortium et al. 2015). Using Snpsift filter v4.3 (Cingolani et al. 2012), we filtered out variants with a quality score less than or equal to 20 or a sample depth of 50 or less. Only biallelic variants were used in our analysis. To filter for somatic variants, we used snpsift filter to remove any variants that had more than 5 alternate allele observations (AO) in the germline sample. Copy number variants were called using the FACETs v0.6.2 R package (Shen and Seshan 2016). Somatic variants found within copy number regions were excluded from the subclone analysis.

To identify subclones within the longitudinal data, we used PyClone-VI v0.1.1 (Gillis and Roth 2020) to cluster the somatic mutations identified in each patient. We ran pyclone-vi fit using the beta-binomial model, 100 restarts, and a maximum of 10 clusters and added each variant’s cluster assignment to its VCF info field. We reconstructed the subclonal architecture of the samples based on this clustering information using SuperSeeker (Qiao et al. 2014). A representation of the subclone structure was then added as a header line to the somatic VCF file for downstream analysis.

### Cell genotyping and assignment

Using the WES VCF file annotated with variant cluster identities and the subclonal structure, cells were genotyped and assigned to subclones using scBayes v1.0.0 (Qiao et al. 2024). Aligned bams containing the long-read scRNA-seq reads for each sample and the VCF file containing the subclone annotations for the given patient were used as input for scBayes. First, we used scGenotype to create a matrix representing the genotype of each cell at all variant positions contained within the VCF file, with one row per variant and one column per cell barcode. scAssign was then used to assign each cell to a subclone based on the presence or absence of subclone-defining mutations in the genotype matrix. We used default parameters for each of these commands.

### Genotype Visualization

To visualize the cell genotypes across all cells in the sample, we developed a method that leverages the genotype matrix and cell assignment from scBayes to produce a genotype matrix plot. Within this scatterplot-like visualization framework, each cell is represented on the X-axis, with the variants of interest along the Y-axis. Cell genotypes at these mutation sites are indicated with a marker: a green marker for cells with only reference alleles at the variant position, a red marker signifying the presence of at least one alternate allele, and no marker present when there is a lack of coverage at the variant position within the given cell. The organization of cells and variants within the plot is done by subclone assignment, enabling a visual depiction of the subclone structure embedded in the genomic data. Within each subclone, cells can be sorted by total read count, number of alternate variants, or assignment quality. By providing a list of disease-driving variants, plots can be tailored to include only cells with these variants to further identify co-occurring driver mutations. This comprehensive visualization approach offers greater insights into the subclonal architecture revealed by long-read scRNA-seq data.

### Coverage metrics

We calculated the percentage of coverage each sequencing read provides to its given transcript by selecting all protein-coding genes annotated in GENCODE v44 (Frankish et al. 2021) for calculation. For each protein-coding gene, we used Pysam to fetch reads that overlapped the genic region, only selecting reads where at least 90% of the read mapped between the gene’s start and end position. Next, we calculated the exonic length of the canonical transcript as annotated by Ensembl (Martin et al. 2023) and determined what percentage of the canonical transcript was covered by a given read. Using these calculations, we determined the percentage of reads that covered X percent of the given transcript for each sample.

In addition, we calculated the fraction of heterozygous germline variants within the WES data that had read coverage within each cell. Using scBayes, we generated a genotype matrix for the germline variants using the scGenotype command. For each variant row in the matrix, we computed the number of cells containing any read that overlapped the variant position, as well as the number of cells with at least one read containing the given variant. The number of cells with variant coverage was divided by the total number of cells present in the sample to determine the percentage of cells covering that variant for plotting.

### Transcriptomic analysis

Seurat v4.3.0.1 (Hao et al. 2021) was used to cluster cells by gene expression and perform a differential gene expression analysis. The sctransform method (Lause et al. 2021) normalized raw gene counts and removed technical variability. The standard steps in the Seurat workflow were then applied, including PCA dimensionality reduction, clustering, and visualization. Dimensions 1 through 20 were used for clustering and UMAP visualization. Subclone assignments were mapped onto cells using the UMAP coordinates of each cell barcode, enabling the identification of subclonal populations within clusters. Inter-subclone differential gene expression analysis was performed on clusters with subclone assignments using the FindMarkers function with the Wilcoxon Rank Sum test. P-values are adjusted using the Bonferroni correction.

### Code availability

The custom scripts and processed datasets generated and/or analyzed in the study are available in the GitHub repository https://github.com/gageblack/BTK-subclones.

## Data access

The single-cell expression data generated in this study have been submitted to the NCBI Gene Expression Omnibus (GEO; https://www.ncbi.nlm.nih.gov/geo/) under accession number GSE259253. The whole exome sequencing data used in this study are available in the NCBI database of Genotypes and Phenotypes (dbGaP; https://www.ncbi.nlm.nih.gov/gap/) under accession number phs003042.v1.p1.

## Acknowledgments

G.S.B., X.H., Y.Q., and G.T.M were supported by National Institutes of Health (NIH) Grant Nos. U24CA209999 and 2U54CA224076-05. G.T.M. is a H.A. and Edna Benning Presidential Endowed Chair. D.M.S is funded by the National Institute of Health, National Cancer Institute R50CA275929. J.A.W. is a clinical scholar of the Leukemia and Lymphoma Society. Research reported in this publication utilized the High-Throughput Genomics and Bioinformatic Analysis Shared Resource at Huntsman Cancer Institute at the University of Utah and was supported by the National Cancer Institute of the National Institutes of Health under Award Number P30 CA042014. The support and resources from the Center for High-Performance Computing at the University of Utah are gratefully acknowledged. The NIH Shared Instrumentation Grant 1S10OD021644-01A1 partially funded the computational resources used for this study. We are grateful to the patients who provided tissue samples for these studies to the OSU Comprehensive Cancer Center Leukemia Tissue Bank Shared Resource (supported by NCI P30 CA016058). We also thank the OSU Leukemia Tissue Bank for assistance with the CLL samples. The content is solely the responsibility of the authors and does not necessarily represent the official views of the National Institutes of Health.

## Author contributions

G.S.B., X.H., Y.Q., D.S., D.M.S., J.A.W., and G.T.M conceived and designed the project. D.M.S. and J.A.W. contributed to the project coordination, clinical oversight, and sample collection and transfer. P.J.M. performed the sample preparation for long-read scRNA-seq. G.S.B performed the data processing of the genomic and long-read scRNA-seq data, performed the subclone and gene-expression analysis, and interpreted the results. X.H., Y.Q., and G.T.M. contributed to the analysis design and assisted in data analysis and the interpretation of results. G.S.B. wrote the manuscript with significant contributions from X.H., Y.Q., and G.T.M. All authors contributed to manuscript editing and refinement.

